# A fast and robust gene knockout method for *Salpingoeca rosetta* informs the genetics of choanoflagellate multicellular development

**DOI:** 10.1101/2024.07.13.603360

**Authors:** Chantal Combredet, Thibaut Brunet

**Affiliations:** Institut Pasteur, Université Paris-Cité, CNRS UMR3691, Evolutionary Cell Biology and Evolution of Morphogenesis Unit, 25-28 rue du docteur Roux, 75015 Paris, France

## Abstract

As the closest living relatives of animals, choanoflagellates offer crucial insights into the evolutionary origin of animals. Notably, certain choanoflagellate species engage in facultative multicellular development that resembles the early stages of embryogenesis. In the past few years, *Salpingoeca rosetta* has emerged as a tractable model for choanoflagellate cell biology and multicellular development, in particular through mutant screens and CRISPR/Cas9-mediated gene knockout (KO). However, existing KO pipelines have variable and sometimes low efficiency, frequently requiring isolation and genotyping of hundreds of clones without guarantee to obtain a KO strain. Here, we present a robust method for gene inactivation in *S. rosetta* that relies on insertion by CRISPR/Cas9 of a single 1.9 kb cassette encoding both a premature termination sequence and an antibiotic resistance gene. We show that this approach allows robust, fast and efficient isolation of KO clones after antibiotic selection. As a proof of principle, we first knocked out all three genes previously reported to regulate *S. rosetta* multicellular development in a published mutant screen (*rosetteless*, *couscous* and *jumble*), and confirmed that all three KOs abolished multicellular development. To showcase the potential of this method for *de novo* characterization of candidate developmental genes, we then inactivated three homologs of genes in the Hippo pathway: *hippo*, *warts* and *yorkie*, which together control cell proliferation and multicellular size in animals. Interestingly, *warts* KO rosettes were consistently about twice as large as their wild-type counterparts, showing our KO pipeline can reveal novel loss-of-function phenotypes of biological interest. Thus, this method has the potential to accelerate choanoflagellate functional genetics.

## Introduction

As the closest living unicellular relatives of animals, choanoflagellates hold the promise to reveal crucial information on animal origins (King, 2004; Leadbeater, 2014; Sebé-Pedrós et al., 2017) (Figure 1A). Beyond their genomic similarities with animals, choanoflagellates display a rich cell and developmental biology, including the ability of some species to differentiate into multiple cell types (Brunet et al., 2021; Dayel et al., 2011; Laundon et al., 2019) and to engage in facultative multicellular development (Brunet et al., 2019; Fairclough et al., 2010; Ros-Rocher et al., 2024).

**Figure 1.**
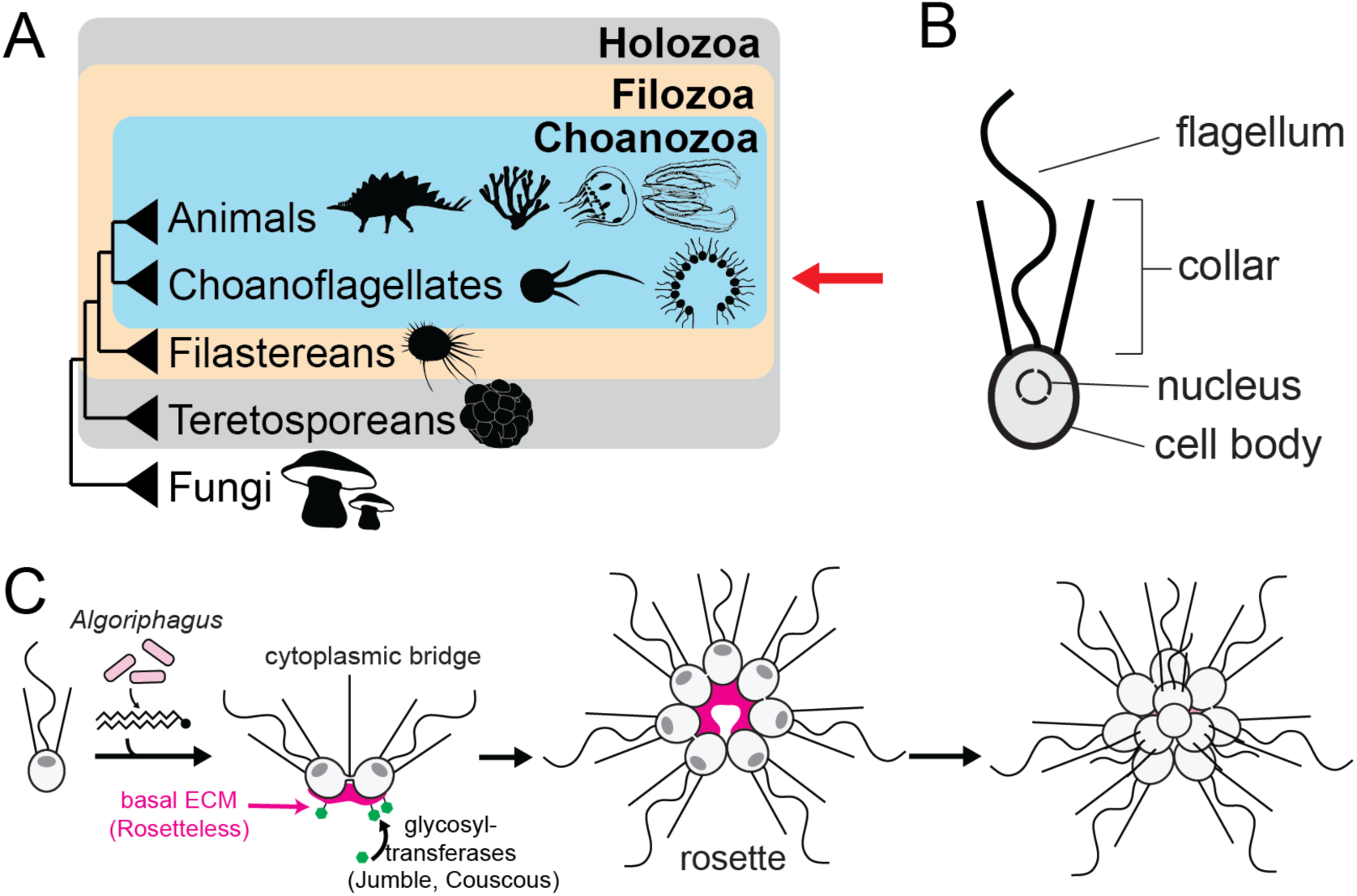
Phylogenetic position and multicellular development of *Salpingoeca rosetta*. (A) Phylogeny of opisthokonts (animals, fungi, and their relatives) showing that choanoflagellates are the sister-group of animals. After (Grau-Bové et al., 2017). (B) Cell morphology of a choanoflagellate, showing the ovoid cell body and the apical flagellum surrounded by a collar of microvilli. After (Brunet and King, 2017; Leadbeater, 2014). (C) Clonal development into spherical rosettes in *Salpingoeca rosetta* (Fairclough et al., 2010). Rosette development is mediated by serial cell division without separation of sister-cells and occurs as a facultative response to lipids secreted by the bacterium *Algoriphagus machipongonensis* (Alegado et al., 2012; Woznica et al., 2016). It relies on secretion of a basal extracellular matrix comprising the C-type lectin Rosetteless (Levin et al., 2014) and glycosylated proteins. Glycosylation is ensured by the glycosyltransferases Jumble and Coucous (Wetzel et al., 2018).

Although choanoflagellates have been known since the mid-19^th^ century (Brunet and King, 2022; Leadbeater, 2014) and their sister-group relationship to animals has been firmly established since the early 2000s (King and Carroll, 2001; King et al., 2008; Ruiz-Trillo et al., 2008), functional genetic tools were lacking. In the past two decades, the choanoflagellate *Salpingoeca rosetta* has emerged as a genetically tractable model organism (Booth and King, 2022). *S. rosetta* develops into spherical multicellular colonies called “rosettes” that resemble the blastula stage of animal embryogenesis (Fairclough et al., 2010). Experimental control was gained over the life history of *S. rosetta*, including its sexual cycle (Levin and King, 2013; Woznica et al., 2017) and its multicellular development (Alegado et al., 2012; Woznica et al., 2016) (Figure 1B,C). A mutant screen (Levin et al., 2014; Wetzel et al., 2018) uncovered three genes necessary for rosette formation: *rosetteless* (*rtls*) (which encodes a lectin that makes part of the extracellular matrix (ECM) at the core of rosettes) as well as *jumble* and *couscous*, that encode predicted glycosyltransferases necessary for proper secretion of the ECM.

Because choanoflagellates are not amenable to microinjection, the development of reverse genetic required the development of methods for transfection with plasmids (Booth et al., 2018; Brunet et al., 2021; Wetzel et al., 2018) and CRISPR/Cas9 ribonucleoproteins (RNPs) (Booth and King, 2020; Coyle et al., 2023; Leon et al., 2024). Previously published approaches for gene knockout (KO) in *S. rosetta* rely on targeted insertion of an 18-basepair translation termination sequence (encoding a stop codon in all 6 possible reading frames). This is achieved by nucleofection (a type of electroporation) of a Cas9:guide RNA complex together with a single-stranded repair template comprising the translation termination sequence flanked by ∼50 bp homology arms. While this approach has allowed successful inactivation of at least 6 distinct genes so far (Booth and King, 2020; Coyle et al., 2023; Leon et al., 2024), a limitation of the existing KO method is its relatively low and variable efficiency. Editing rates ranging from 0.3% to 16.5% in published successful KOs (Booth and King, 2020; Leon et al., 2024), often requiring isolation and genotyping of hundreds of clones to potentially isolate a KO strain. Because choanoflagellates are aquatic, clonal isolation is labor-intensive and cannot be done on solid media. Clonal isolation and genotyping require about a month and entail some uncertainty: indeed, in some experiments, no KO strain could be obtained after isolating and genotyping hundreds or thousands of clones, even for non-essential loci (Booth and King, 2020; Coyle et al., 2023). Although efficiency was increased in certain experiments by co-editing an independent locus (*rpl36a*) to confer resistance to cycloheximide, KO efficiency after cycloheximide selection was still as low as 0.6% (Coyle et al., 2023). Moreover, co-editing for cycloheximide resistance actually decreased KO efficiency at certain loci (such as *cRFXa* (Coyle et al., 2023)), suggesting it might not be a universally applicable solution. Thus, a highly efficient method for gene inactivation has been lacking.

Here, we establish a novel KO method for *S. rosetta* relying on insertion of a premature termination site followed by an antibiotic resistance gene in the coding sequence of the target gene. This allows direct production and selection of KO cells with a single editing event. The method relies on transfection of ∼2 kb double-stranded repair templates produced by PCR, which are affordable and straightforward to generate. We summarize below the establishment of this method and report on the phenotypes of 6 novel KO strains generated as a proof of principle, three of which recapitulate known mutants while the three others inactivate previously uncharacterized candidate genes for multicellular development. We could knock out all genes we attempted and the frequency of KO among puromycin-resistant clones ranged from 40% to 100% – an increase of two orders of magnitude in efficiency compared to earlier methods. We show that this method has the potential to accelerate future functional studies in *S. rosetta*.

## Results and Discussion

### Design and production of repair templates for insertion of a termination/resistance cassette

To increase the efficiency and reliability of genome editing in *S. rosetta*, we set out to disrupt the open reading frame of genes of interest by insertion of an antibiotic resistance cassette directly in the target locus. We reasoned that, if successful, this would allow antibiotic-mediated selection of KO cells, similar to established pipelines in other species including yeast (Wach et al., 1994), mice (Hall et al., 2009), and *Capsaspora owczarzaki* (a close relative of choanoflagellates and animals) (Phillips et al., 2022). An earlier study had shown that *S. rosetta* readily repaired Cas9-induced double-stranded breaks by homologous recombination (homology-dependent repair, or HDR) between the genome and co-transfected repair templates (Booth and King, 2020). We thus designed repair templates with the following properties: (1) short (50, 80 or 155 bp) homology arms to the target locus (flanking both sides of the Cas9 cleavage site); (2) a premature termination sequence encoding stop codons in all 6 possible reading frames; (3) a puromycin resistance cassette, including an open reading frame encoding the *pac* resistance gene preceded by a strong *S. rosetta* promoter (the EFL promoter or pEFL) and followed by the 3’UTR of a highly expressed gene (Actin-3’UTR; the full resistance cassette will be referred to as pEFL-pac-Act3’) (Fig. 2A). Although commonly used protocols for *S. rosetta* genome editing rely on single-stranded repair templates, published data indicate that *S. rosetta* can also successfully incorporate double-stranded repair templates by HDR, albeit with lower efficiency (Booth and King, 2020). We decided to use double-stranded templates, as they can be easily and affordably produced by PCR. A published plasmid for *S. rosetta* puromycin resistance (Brunet et al., 2021) was used as a PCR template, with custom primers including premature termination sites and homology arms (Fig. S1). We included premature termination sites at both ends of replair templates (Fig. 2A, Fig. S1) to achieve KO regardless of the orientation of the insert relative to the reading frame of the target gene.

**Figure 2.**
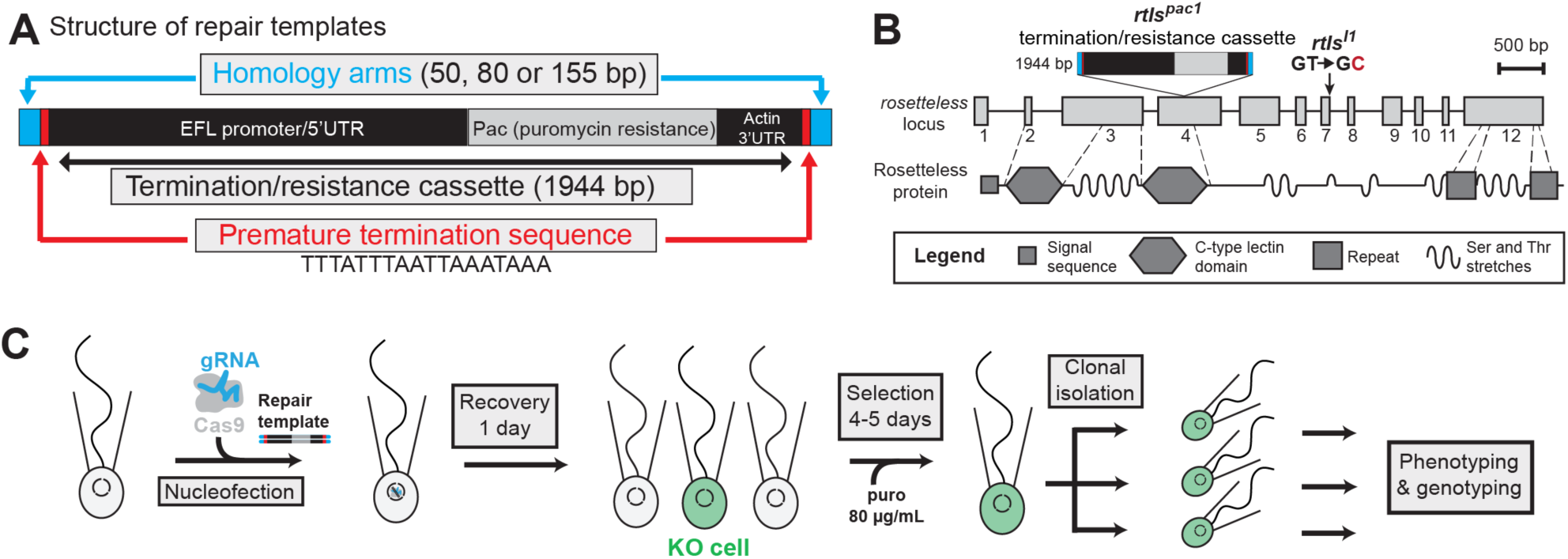
Principle of *rosetteless* knock-out by insertion of a puromycin resistance cassette. (A) Architecture of transfected repair templates, comprising homology arms flanking the CRISPR/Cas9 cleavage site, premature termination sequences on both ends, and a termination/resistance cassette comprising the Pac1 puromycin resistance gene under the control of a strong *S. rosetta* promoter (that of the *efl* gene) and with the 3’ UTR of another highly expressed gene (that of *S. rosetta actin*). Repair templates are double-stranded and generated by PCR (see Fig. S1 and Material and Methods). (B) Structure of the *rosetteless* locus and the Rosetteless protein, together with the originally isolated loss-of-function mutation *rtls^l1^* (loss of a splice donor site between exons 7 and 8). A previously published gRNA (Booth and King, 2020) targeting exon 4 was used to introduce the termination/resistance cassette (or pac-stop cassette) and prematurely interrupt translation of the *rosetteless* gene. (C) Knockout pipeline. A ribonucleoprotein (Cas9:gRNA) complex is transfected together with the double-stranded repair template, and clones are isolated by puromycin (“puro”) selection followed by limiting dilution. Modified from (Booth and King, 2020).

### Targeted insertion of the termination/resistance cassette allows fast, robust and specific knockout

As a first test case, we aimed to disrupt the *rosetteless* gene (PTSG_03555), as its loss-of-function phenotype is well-described and visually easy to score (Booth and King, 2020; Levin et al., 2014). We transfected a published guide RNA against *rtls* complexed with recombinantly produced Cas9 nuclease (Booth and King, 2020), together with repair templates comprising the termination/resistance cassette. We tested three different lengths of homology arms (50 bp, 80 bp, and 155 bp): although 50 bp homology arms have been successfully used in the past to insert short sequences (Booth and King, 2020), we reasoned that longer homology arms might facilitate the insertion of longer cassettes. Finally, we included two negative controls: (1) cells transfected with the Cas9/guide RNA ribonucleoprotein complex (RNP) but without repair templates; (2) cells transfected with repair templates but without RNP.

Cells were transfected using a published protocol relying on the Lonza 4D nucleofector, treated with 80 µg/mL puromycin (Fig. 2C), and monitored for the appearance of resistant cells (which we recognized by their ability to proliferate, and thus to develop into clonal chains of cells, under puromycin selection). After 4 to 5 days, resistant cells could readily be observed in each of the six wells co-transfected with RNP and one of the three types of repair templates. No resistant cells were observed in negative control wells that had undergone nucleofection without repair templates. For each type of repair template tested, we isolated 13 to 45 resistant clones by limiting dilution and proceeded to induce rosette development by treating clones with *Algoriphagus* (Table 1). This assay allows straightforward identification of *rtls* loss-of-function mutants based on their inability to develop into rosettes (Fig. 3A-C).

**Figure 3.**
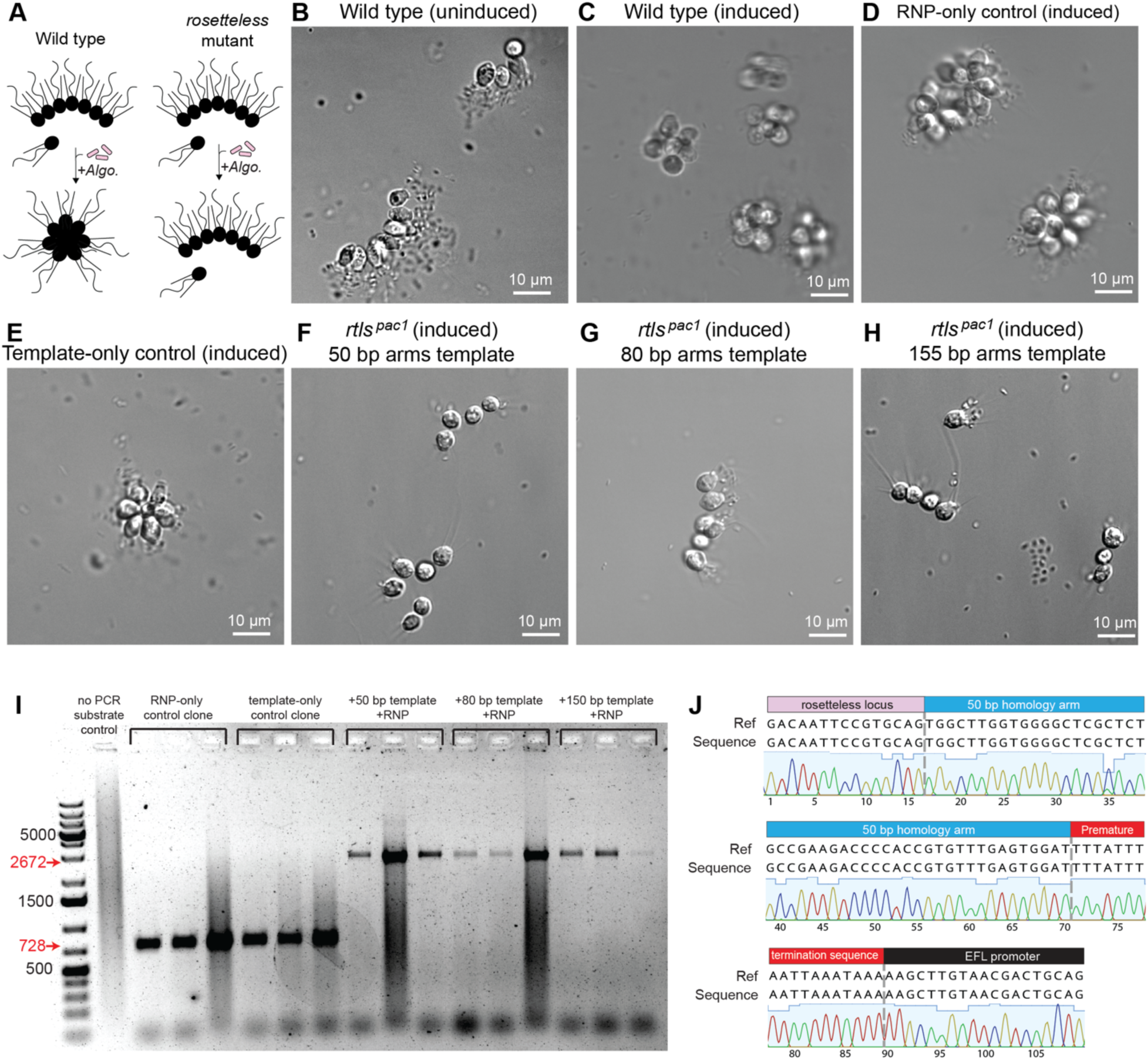
Phenotype and genotype of *rtls^pac1^* knock-out clones. (A) Described phenotype of *rosetteless* loss-of-function mutants. In the absence of *Algoriphagus* and in co-culture with the standard food bacterium *Echinicola pacifica*, *S. rosetta* develops into linear chain colonies. Upon addition of *Algoriphagus*, wild type cells develop into spherical rosette colonies instead, while *rosetteless* loss-of-function mutants remain in chains. (B-H) Phenotypes of knockout clones generated by insertion of the termination/resistance cassette in the *rosetteless* locus. All puromycin-resistant clones generated by transfection of RNP with repair templates (with 50, 80 or 155 bp homology arms) displayed the *rosetteless* KO phenotype. Wild-type cells, RNP-only control clones, and template-only control clones, all displayed the wild-type phenotype. Note that template-only control clones were resistant to puromycin. See Table 1 for statistics. (I) Genotyping PCR confirmed insertion of the termination resistance/cassette in the *rosetteless* locus in 9 co-transfected clones displaying the *rosetteless* loss-of-function phenotypes (3 for each length of homology arm). Wild-type PCR product has a length of 728 bp, while the *rtls^pac1^* PCR product has a predicted length of 2,672 bp. (J) Sanger sequencing of PCR products confirms insertion of the termination/resistance cassette in the *rtls* locus. The clone sequenced in this example was produced by transfecting 50-bp homology arms repair template. Ref: predicted reference sequence of the *rtls^pac1^*locus under the hypothesis of a specific insertion of the termination/resistance cassette. Seq: sequence of the PCR amplicons as determined by Sanger sequencing.

**Table 1.** Number and phenotypes of clones isolated after transfection of *rtls^pac1^* repair constructs and RNPs. The table reports the results of two biological replicates (independent transfection experiments: Exp 1 and Exp 2). All clones were isolated by limiting dilution and induced for rosette development by addition of *Algoriphagus machipongonensis*. Note that in the no-repair template controls (transfected with RNP only), no puromycin resistant cells were observed in either replicate. In that condition only, clones were isolated and characterized without puromycin selection.

| Electroporated construct | Puromycin selection |  | Clones forming rosettes |  | Clones not forming rosettes |  |
| --- | --- | --- | --- | --- | --- | --- |
|  | Exp 1 | Exp 2 | Exp 1 | Exp 2 | Exp 1 | Exp 2 |
| RNP + *rtls<sup>pac1</sup>* with 50 bp homology arms | Yes | Yes | 0 | 0 | 29 | 20 |
| RNP + *rtls<sup>pac1</sup>* with 80 bp homology arms | Yes | Yes | 0 | 0 | 35 | 21 |
| RNP + *rtls<sup>pac1</sup>* with 155 bp homology arms | Yes | Yes | 0 | 0 | 45 | 37 |
| *rtls<sup>pac1</sup>* with 50 bp homology arms (template-only control) | Yes | Yes | 39 | 5 | 0 | 0 |
| RNP (RNP-only control) | No (no resistant cells observed) | No (no resistant cells observed) | 13 | 29 | 0 | 0 |

Strikingly, all clones co-transfected with RNP and repair templates displayed the *rtls* mutant phenotype (187/187 clones across two biological replicates and three tested lengths of homology arms; Table 1; Fig. 3F-H), suggesting that the repair templates had been integrated at the target locus and had disrupted the *rtls* gene with high efficiency. Wild-type control cells treated in parallel with *Algoriphagus* developed into canonical rosettes. To confirm *rtls* KO, we genotyped three clones for each length of homology arms by PCR (nine clones in total). 9/9 clones showed evidence of a ∼2 kb insertion in the *rtls* locus, suggesting successful integration of the termination/resistance cassette (Fig. 3I) which was confirmed by Sanger sequencing (Fig. 3J). We named the corresponding loss-of-function *rtls* allele *rtls^pac1^* (after the *pac* gene, conferring resistance to puromycin). We also assessed specificity of the termination/resistance cassette insertion by whole-genome sequencing of five *rtls^pac1^*clones generated with 50-bp repair templates and detected a unique insertion of the termination/cassette at the *rtls* locus in 5/5 clones (Fig. S2; Table S1; Table S2). Finally, quantification of cell growth showed that *rtls^pac1^* clones proliferated about 9% slower than wild-type cells (doubling time 8.9 ± 0 .3 versus 8.1 ± 0.2 hours, respectively; Fig. S3), similar to the previously characterized loss-of-function alleles, *rtls^PTS1^* (generated by CRISPR/Cas9-mediated editing) and *rtls^tl1^* (generated by random mutagenesis) (Booth and King, 2020). This suggested that the energetic cost of expressing the puromycin resistance cassette did not measurably slow down cell proliferation. Taken together, these observations support high specificity and efficiency of selection-mediated *rtls* KO.

### Both RNP and repair templates are necessary for KO

In parallel, we produced and characterized negative control clones transfected either with RNP only (without repair templates) or with repair templates only (without RNP). No resistant cells were obtained in the RNP-only control condition, but unexpectedly, resistant cells were observed in template-only control wells. We set out to test whether *rtls* KO had been achieved in these cells in the absence of RNP – perhaps by integration of the repair template in the absence of Cas9-induced DNA breaks. We isolated 13 puromycin-resistant template-only control clones and treated them with *Algoriphagus.* All developed into rosettes (13/13; Fig. 3E), as did (non-resistant) RNP-only control clones isolated without puromycin selection (13/13, Fig. 3D). This suggested that the *rtls* locus had not been disrupted in template-only controls. Instead, the puromycin resistance cassette had presumably been integrated at another locus or was maintained as an episomal element outside the genome.

To determine whether (and where) the termination/resistance cassette was integrated, we sequenced the whole genome of two puromycin-resistant template-only control clones. To our surprise, we found no evidence of integration. Rather, reads indicated that the repair template had undergone circularization by recombination between its left and right homology arms to form a 2-kb episomal circular element in both clones (Fig. S4; Table S1; Table S2). Additionally, one of the two clones only showed evidence of recombination between the repair template and the pUC19 plasmid used as carrier during transfection, forming chimeric plasmids present alongside the circularized repair template. Both the circularized repair template and the chimeric pUC19/template plasmid (when present) were estimated to be present in 4 to 7 copies per genome (based on relative read coverage). This suggested that *S. rosetta* had maintained, and presumably replicated, these episomal elements over the 21 days of our experiment, suggesting that an unidentified sequence within these constructs had acted as an origin of replication. The fact that no such episomal elements were detected in the five *rtls^pac1^* clones we sequenced suggests that such events of circularization/recombination are significantly rarer than targeted insertion after RNP co-transfection. Taken together, these observations suggest that efficient KO requires co-transfection of both RNP and repair templates, and cannot be achieved by transfecting repair templates alone.

### KO of *couscous* and *jumble* confirms the role of glycosyltransferases in rosette development

We then set out to test the versatility of the method by targeting two other genes previously proposed to be required for rosette development, *jumble* (PTSG_00436) and *couscous* (PTSG_07368), which encode predicted glycosyltransferases (Wetzel et al., 2018). We designed guide RNAs targeting the coding sequence of each locus, both located 5’ of the published loss-of-function mutations (Fig. 4A,C). We co-transfected Cas9/guide RNA complexes with repair templates comprising the *pac* locus flanked by 50 bp homology arms. After puromycin selection, we isolated and genotyped 5 resistant clones per target gene. 2/5 and 5/5 clones were found to be knockouts for *jumble* and *couscous*, respectively (Fig. 4B,D); we refer to the corresponding knockout alleles as *couscous^pac1^*and *jumble^pac1^*. All knockout clones phenocopied the published *couscous* and *jumble* mutant phenotypes characterized by large, irregular clumps of cells (Fig. 4E-G) and by an inability to develop into rosettes upon *Algoriphagus* treatment (with irregular aggregates persisting instead (Fig. 4H-J)). In addition to their size and shape, the clumps formed by the originally described *couscous* and *jumble* mutants differed from rosettes by their physical properties: clumps readily dissociated upon vortexing, while rosettes were unaffected (Wetzel et al., 2018). In line with published observations of these mutants, we found that clumps formed by *Algoriphagus*-treated *couscous^pac1^* and *jumble^pac1^* clones dissociated into single cells in a matter of seconds upon vortexing (Fig. 4L,M). Wild-type rosettes, on the other hand, remained cohesive, as reported previously (Wetzel et al., 2018) (Fig 4K). Taken together, these results demonstrate the versatility of our knockout protocol and provide independent confirmation for the necessity of these predicted glycosyltransferases for rosette development in *S. rosetta*.

**Figure 4.**
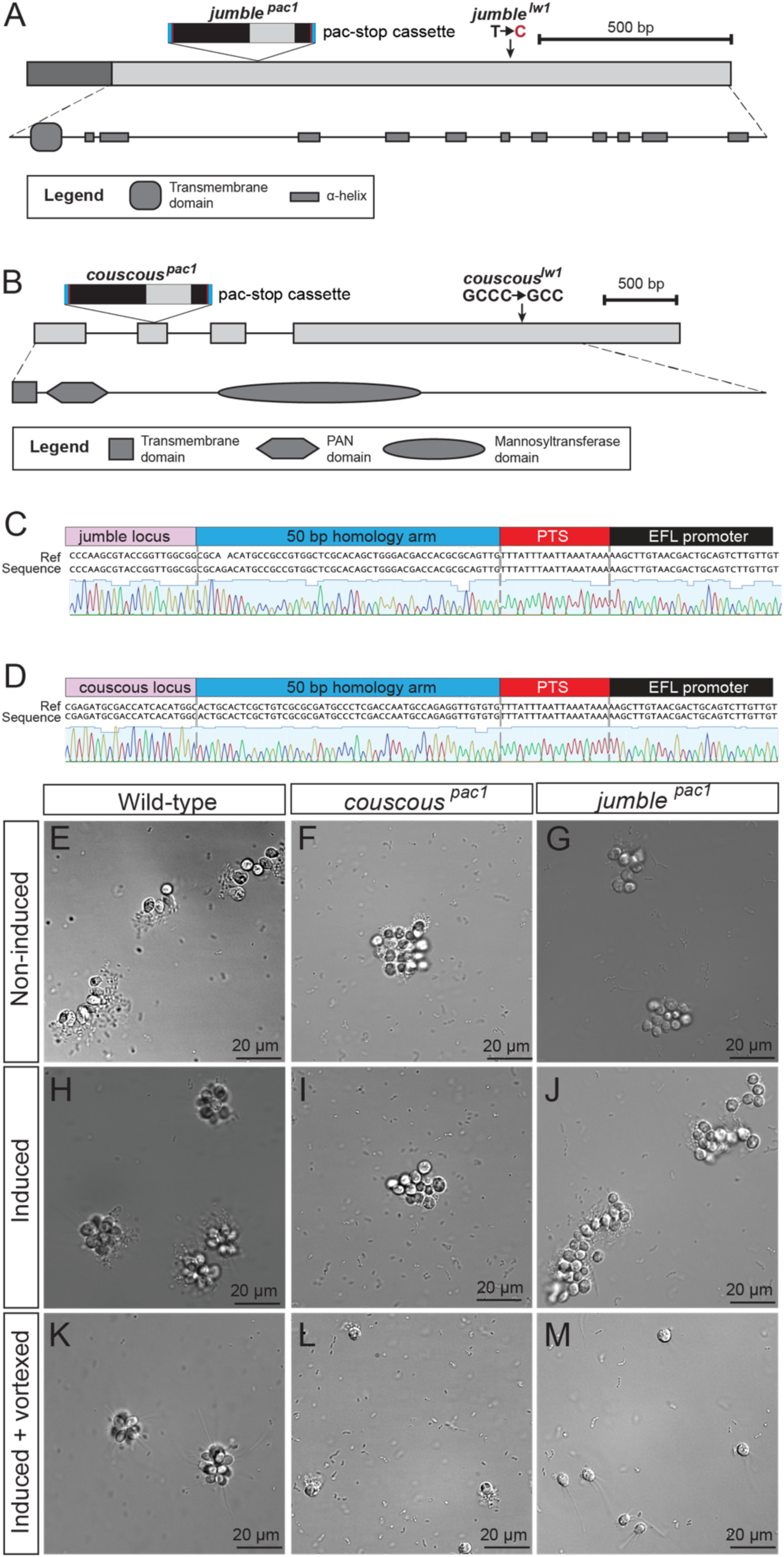
Knockout of *jumble* and *couscous* abolishes rosette development and promotes spurious cell clumping. (A,B) Structure of the *jumble* and *couscous* locus and of the Jumble and Couscous proteins, respectively. Loss-of-function alleles mapped in a published mutant screen are indicated (*jumble^lw1^* and *couscous^lw1^*) as well as insertion sites for the termination-resistance cassette for the newly generated loss-of-function alleles *jumble^pac1^* and *couscous^pac1^*. (C,D) Sanger sequencing results of *jumble^pac1^* and *couscous^pac1^*knockout clones. (D) *jumble^pac1^* and *couscous^pac1^*phenocopy *jumble^lw1^* and *couscous^lw1^*. Both knockout strains form spurious cell aggregates instead of chains and rosettes when they are cultured respectively in the absence or in the presence of *Algoriphagus*. Aggregates differ from rosettes by their irregular size and shape, as well as by the fact that they dissociate upon vortexing.

### Inactivation of Hippo pathway genes reveals a role for Warts kinase in setting rosette size

Finally, we set out to test the potential of our pipeline to provide new insights into candidate *S. rosetta* developmental genes. We focused on the Hippo pathway, which controls proliferation and multicellular cell size in animals (Fig. 5A) (Meng et al., 2016; Pan, 2022). Interestingly, three core components of the Hippo pathway (the transcription factor Yorkie, that stimulates proliferation, and the kinases Hippo and Warts, that inactivate Yorkie by phosphorylation) are present in choanoflagellates, as well as in filastereans (the second closest living relatives of animals; Fig. 1A) (Phillips et al., 2024; Sebé-Pedrós et al., 2012). In the filasterean *Capsaspora*, functional studies indicate that Hippo, Warts and Yorkie regulate cell contractility, and thereby the shape of multicellular aggregates (Phillips and Pan, 2023; Phillips et al., 2022), without clearly impacting proliferation rate or multicellular size. This has given rise to the hypothesis that the function of the Hippo pathway in the control of growth had evolved after the divergence of filastereans – either in stem choanozoans, or within metazoans (Phillips et al., 2024).

**Figure 5.**
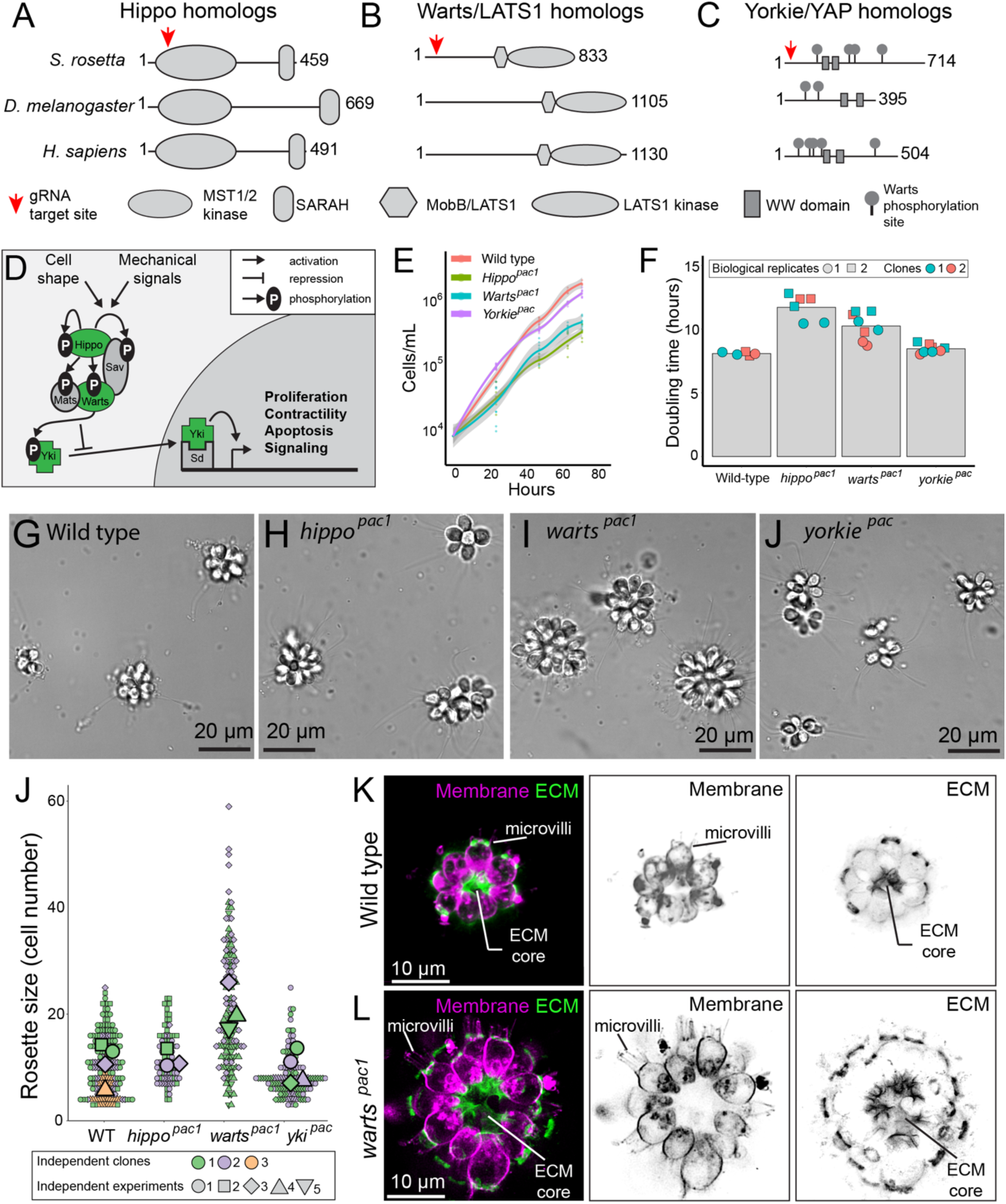
Knockout of *S. rosetta* homologs of Hippo pathway genes suggests a role for Warts kinase in controlling rosette size. (A-C) Domain architecture of Hippo pathway proteins in *Homo sapiens* and *Drosophila melanogaster* and of their orthologs in *S. rosetta*, identified by (Sebé-Pedrós et al., 2012). Sequences were retrieved from UniProt (The UniProt Consortium et al., 2023) and domain architectures predicted by NCBI CD Search (Marchler-Bauer and Bryant, 2004). (D) Architecture of the Hippo pathway, after (Bardet, 2009). Proteins whose homologs were targeted for KO are highlighted in green. (E) Growth curves of wild type and Hippo pathway knockout cells. *yorkie^pac1^*and wild type cells grew at a similar rate but *hippo^pac1^* and *warts^pac1^* grew slower. Curves represent results of 1/2 independent biological replicate (see Fig. S6 for the second biological replicate). (F) Doubling times calculated for each genotype. *hippo^pac1^* and *warts^pac1^*had significantly longer doubling times than wild type cells (*p=*0.2% and 0.4% by the Mann-Whitney test, respectively). Data represented comprise both biological replicates (panel E and Fig. S6). (G-I) Phenotypes of wild type *S. rosetta* compared to *hippo^pac1^*, *warts^pac1^* and *yorkie^pac^* KO strains after induction with *Algoriphagus*. All genotypes readily developed into rosettes but *warts^pac1^* rosettes were conspicuously larger. Two independent clones were analyzed for each knockout genotype. (J) *warts^pac1^*rosettes displayed a significantly higher number of cells per rosettes (ANOVA *p*=0.7%). All other conditions were statistically identical. (K, L) Both wild-type and *warts^pac1^* rosettes display a central ECM core, which appears larger and more branched in *warts^pac1^*rosettes.

We designed gRNAs targeting the beginning of the coding sequence of the *S. rosetta* homologs of Hippo (PTSG_10780), Warts (PTSG_04961) and Yorkie (PTSG_06057) (Fig. 5B-E) and isolated two knockout clones for each gene by insertion of the puromycin resistance cassette (with a respective success rate of 2/5, 2/5, and 5/5 KOs among genotyped clones; Fig. S5). For *yorkie* only, both KO clones had subtly different sequences from each other (Fig. S5) and the two alleles were thus named *yorkie^pac1^* and *yorkie^pac2^*, collectively referred to as *yorkie^pac^*. Growth curves indicated that *yorkie^pac^* KO cells proliferated at a similar rate to the wild type (doubling time 8.5 ± 0.3 and 8.1 ± 0.1 hours, respectively). On the other hand, *hippo^pac1^* and *warts ^pac1^* KO clones proliferated markedly slower (doubling time 11.7±1.0 and 10.2±1.1 hours, respectively) (Fig. 5F). After treatment with *Algoriphagus*, all six KO clones reliably developed into rosettes, as did wild-type cells. Strikingly, *warts ^pac1^* cells grew into giant rosettes containing about twice as many cells as wild-type ones (21.1 ± 4.4 versus 10.9 ± 3.8 cells per rosette, respectively) (Fig. 5F-J). *warts^pac1^* rosettes contained as many as 60 cells, while their wild type counterparts did not exceed 25 cells (Fig. 5J), consistent with earlier quantifications of wild type rosette size (Larson et al., 2020). The size of *hippo^pac1^* and *yorkie^pac^* rosettes did not significantly differ from wild type (11.6 ± 1.7 cells and 9.8 ± 3.1 cells, respectively) (Fig. 5F-J).

In wild-type *S. rosetta*, rosette size and shape is thought to be set by adhesion of cells to a common core of self-secreted extracellular matrix (Larson et al., 2020). To visualize the structure of giant *warts ^pac1^* rosettes, we performed confocal imaging after staining the ECM with a fluorescent lectin (jacalin-fluorescein) and cell outlines with the lipid dye FM 4-64. Like the wild type, *warts ^pac1^* rosettes were structured as monolayers of cells surrounding a core of ECM (Fig. 5K,L). However, that ECM core appeared larger and with a more conspicuously branched geometry in *warts ^pac1^* rosettes (Fig. 5K,L). Interestingly, the branched ECM of *warts ^pac1^*rosettes evoked that of the choanoflagellate species *Barroeca monosierra*, a close relative of *S. rosetta* that naturally develops into much larger rosettes (2.5 times as large in diameter) (Hake et al., 2021).

Overall, these data suggest that components of the Hippo pathway influence both cell division rate and multicellular size in *S. rosetta* – as they do in animals. As in animals, *warts* knockout caused hypertrophic growth of *S. rosetta* rosettes. However, *hippo* and *warts* loss-of-function reduced the rate of cell proliferation in *S. rosetta*, which is the opposite of their animal loss-of-function phenotype. Deeper elucidation of the cellular and molecular mechanisms underlying these phenotypes, and reconstitution of the evolution of the Hippo pathway, will require dedicated functional studies. These observations pinpoint *warts* as a genetic regulator of *S. rosetta* rosette size, and exemplify the ability of our KO pipeline to inform the developmental biology of *S. rosetta*.

## Conclusion

Here, we have reported a robust method for gene inactivation in *S. rosetta*. We could knock out all of the six genes we targeted, with the frequency of KO among puromycin-resistant clones ranging from 40% to 100%. Besides its efficiency, our method brings about a significant reduction in duration of the experiments and sequencing costs compared to earlier pipelines. We thus think it has the potential accelerate the functional characterization of *S. rosetta* and enriches the growing molecular toolkit available for diverse unicellular relatives of animals (Kożyczkowska et al., 2021; Parra-Acero et al., 2018; Phillips et al., 2022; Suga and Ruiz-Trillo, 2013; Woznica et al., 2021) The present method can still be improved in some respects. Although the resistant KO clones we sequenced all displayed a single specific insertion of the resistance cassette, we sometimes also isolated resistant clones that were not KO. One factor might be occasional circularization of repair cassettes followed by their maintenance as episomal plasmid-like elements, as observed in transfection of repair templates without RNP. How these elements are replicated and transmitted during cell division, and notably the nature of the sequence acting as an origin of replication, should be the object of future studies.

Beyond the specific application to gene inactivation, our study shows that *S. rosetta* can incorporate large, ∼2 kb inserts by homologous recombination at a precise target site. This represents an increase of 2 orders of magnitude in size compared to previous insertions in *S. rosetta* and might open the door to other applications, such as knock-in of fluorescent tags fused to proteins of interest (Paix et al., 2015; Paix et al., 2017; Paix et al., 2023; Seleit et al., 2021). Finally, a current limitation of this method is that is only allows selection-mediated KO of a single gene. Multiple KOs will be facilitated when other selectable markers (beyond puromycin resistance) are identified.

Future efforts at improving editing efficiency will hopefully yield even more robust protocols, potentially achieving KO with efficiency and specificity close to 100%. We anticipate this will pave the way for large-scale efforts such as genome-wide knockout screens.

## Acknowledgements

We thank David Booth, Alain Garcia de las Bayonas, Nicole King, Juliette Mathieu, Núria Ros-Rocher, and all members of the ECB lab at the Institut Pasteur for comments on the manuscript and feedback during the project. Work in the ECB lab is supported by the Institut Pasteur (G5 package), the ERC Starting Grant EvoMorphoCell (Grant agreement ID: 101040745), the Bert L. and N. Kuggie Vallee Foundation, the ANR-23-CE13-0031, and the CNRS (UMR 3691). Funded by the European Union. Views and opinions expressed are however those of the author(s) only and do not necessarily reflect those of the European Union or the European Research Council. Neither the European Union nor the granting authority can be held responsible for them.

## Material and Methods

All centrifugation steps were performed either on an Eppendorf 5415R tabletop microcentrifuge (for volumes up to 2 mL) or on a FisherBrand GT2R expert tabletop centrifuge (for volumes between 2 mL and 50 mL).

### Choanoflagellate cultures

*Salpingoeca rosetta* was maintained in monoxenic co-culture with the bacterium *Echinocola pacifica* (co-culture “SrEpac”; American Type Culture Collection ATCC PRA-390 and (Levin and King, 2013)). Cells were grown in 5% (v/v) SeaWater Complete medium (SWC) in Artificial SeaWater (ASW) prepared from Instant Ocean powder (Aquarius System) as established in (Levin and King, 2013) and following modifications in (Ros-Rocher et al., 2024). At each passage, cultures were supplemented with 1% (v/v) of a 10 mg *E. pacifica* pellet (hereafter “*E. pacifica* food pellet”; (Booth et al., 2018)) resuspended in 1 mL ASW to ensure regular supply of food bacteria. Cultures were passaged up to 20 times before being re-established from a cryogenically preserved stock (King et al., 2009). Cultures were maintained in a 25°C incubator (Memmert IPP410ecoplus) under a 12-hour light-dark cycle. For long-term storage of wild type and mutant strains, cryogenic preservation was performed as in (King et al., 2009), using 10% (v/v) glycerol instead of 10% (v/v) DMSO.

### Design and preparation of guide RNAs

Guide RNAs (gRNA) were designed and prepared as in (Booth and King, 2020). In brief, guides RNA were prepared by annealing a gene-specific CRISPR RNA (crRNA) with an invariant trans-activating CRISPR RNA (trRNA), both purchased from Integrated DNA Technologies (IDT DNA, Coralville, IA, USA). crRNAs were designed on the EuPaGDT platform http://grna.ctegd.uga.edu/ (Peng and Tarleton, 2015) to target sites (aiming to bind as close as possible to the 5’ end of the coding sequence). Whenever possible, we followed some constraints on the base pairs surrounding the PAM site (5’-HNNRR<u>VGG</u>H-3’ or ‘strict PAM sequence’ where the standard PAM is underlined), in order to maximize binding to Cas9 and to the target site. For some genes (*hippo*, *yorkie* and *warts*), we could not find crRNAs close to the 5’ end following the strict PAM sequence, and used a minimal PAM instead (5’-NGG-3’). Possible crRNAs were selected based on proximity to the translation start site (to achieve maximal truncation of the target protein) and further screened based on predicted secondary structure. Only crRNAs with a folding energy ΔG > −2.0 kcal/mol (and preferentially > −1.5 kcal/mol) were considered. Folding energies were calculated on the RNAfold Web Server http://rna.tbi.univie.ac.at/cgi-bin/RNAWebSuite/RNAfold.cgi (ViennaRNA Web Services (Gruber et al., 2015)). Sequences of crRNAs used are present in Supplementary File 1. crRNAs and a trRNA stock were resuspended and annealed *in vitro* as in (Booth and King, 2020).

### Design and preparation of repair templates

Repair templates were produced by PCR. A published plasmid encoding the pEFL-pac-5’Act cassette was used as a template (Addgene ID pMS18, catalog number #225681). Custom primers were designed to add an 18-nucleotide premature termination cassette (5’-TTTATTTAATTAAATAAA-3’) and 50 bp, 80 bp, or 155 bp homology arms. Sequences of all primers used are present in Supplementary File 1. Primers were purchased from Eurofins Genomics (for 50 bp or 80 bp homology arms) or from IDT DNA (for 155 bp homology arms). PCR reactions were set up in 100 µL as follows using Q5 high fidelity DNA polymerase (New England Biolabs M0491S) following provider specifications, with 100 ng plasmid as template, 200 µM dNTP (ThermoScientific R1121), 0.5 µM of each primer, 0.02 U/µL Q5 polymerase.

The following PCR program was run on a PCRmax AlphaCycler 2 with 96-well blocks: 30” 98°C; 40X (10” 98°C; 30” 68°C; 1’ 72°C); 2’ 72°C; hold 10°C. Size of a PCR products (expected: 2 kb) was visually checked by running 1:10 of the PCR reaction product on a 1% (w/v) agarose (Invitrogen 1015) gel, run in Tris acetate EDTA (TAE, Euromedex EU0201) and visualized with a Safe Imager 2.0 Blue-Light Transilluminator (Invitrogen G6600EU). PCR products were purified using a PCR purification kit (Macherey NucleoSpin Gel and PCR cleanup 740609.50) and the final product was eluted with 17 µL water. 2 µL were used to measure DNA concentration using a NanoDrop spectrophotometer (LabTech). The remaining 15 µL were reconcentrated in 2-3 µL by evaporation for 1 hour at 55°C and then used as repair template for nucleofection in the following quantity:

|  |  |
| --- | --- |
| Rosetteless (50 bp homology) | 6 μg (experiment 1); 6.9 μg (experiment 2) |
| Rosetteless (80 bp homology) | 4.5 μg (experiment 1); 4.6 μg (experiment 2) |
| Rosetteless (155 bp homology) | 2 μg (experiment 1); 5.6 μg (experiment 2) |
| Couscous | 7.1 µg |
| Jumble | 2.4 µg |

### Nucleofection

The nucleofection protocol was modified from (Booth and King, 2020). Main steps are summarized below.

### Setting up *S. rosetta* cultures

48 hours before the experiment, 120 mL of SrEpac culture were set up at 8,000 cells/mL in a 25°C incubator (Memmert IPP410ecoplus) in which humidity was maintained by evaporation of a tank of distilled water.

### Preparing gRNA and RNP

gRNAs were prepared by hybridizing crRNA with synthetic transactivating CRISPR RNA (trRNA). gRNA complexes were produced by suspending crRNA and trRNA in duplex buffer (30 mM HEPES-KOH pH 7.5, 100 mM potassium acetate) to a final concentration of 200 µM each. crRNA and trRNA were then mixed 1:1, resulting in a final concentration of 100 µM each in duplex buffer. The mix was incubated at 95°C on a heating block for 5 minutes, and then let to gently cool off to room temperature (at least 2 hours) to support annealing. gRNA was either stored at −20°C or used immediately. gRNAs were then complexed with Cas9 to form RNPs. For each transfection reaction, 2 µL of 20 µM SpCas9 (New England Biolabs M0646T) were gently mixed with 2 µL of 100 µM gRNA, incubated for 1 hour at room temperature to assemble into an RNP (in parallel with repair template reconcentration at 55°C; see “Design and preparation of repair templates”). RNPs were kept on ice before use.

### Concentration of *S. rosetta* cultures and reduction of bacterial density

48 hours before the experiment, SrEpac cultures were seeded at 8,000 cells/mL in 120 mL. The full volume of SrEpac culture were harvested by repeated rounds of centrifugation (to reduce bacterial density) in 50 mL tubes using the following protocol: 5’ 2,000g; pellets were resuspended in ASW and consolidated in a single 50 mL Falcon tube; 5’ 2,000 g; 5’ 2,2000 g. Supernatant was carefully removed and pellet was resuspended in 400 µL ASW (hereby referred to as “washed cells”). Cell concentration was quantified using a LUNA-II^TM^ automated cell counter (LogosBiosystems) after immobilizing cells with 0.16% paraformaldehyde (by plating 10 µL of the following suspension: 2 µL of washed cells + 196 µL ASW + 2 µL 16% paraformaldehyde, Electron Microscopy Sciences 15714). Cells were then diluted down to 5E7 cells/mL and split into 100 µL aliquots. Each aliquot was sufficient for 12 nucleofection reactions.

### Glycocalyx digestion

Each 100 µL aliquot of washed cell was centrifuged 5’ at 5,000 g. The pellet was resuspended in 100 µL of pre-treatment solution.

To prepare the pre-treatment solution, 5 µL commercial papain solution (Sigma Aldrich P3125-100MG) were first diluted in 45 µL papain dilution buffer (50 mM HEPES-KOH pH. 7.5, 200 mM NaCl, 20% (v/v) glycerol, 10 mM L-cysteine; 0.22 µM-filtered and stored at −20°C). 4 µL of the resulting diluted papain solution were then mixed with 400 µL priming buffer (40 mM HEPES-KOH pH 7.5, 34 mM lithium citrate, 50 mM L-cysteine, 15% PEG 8000; 0.22 µM-filtered and stored at −20°C). This results in a final ‘pre-treatment solution’ containing 1 µM papain in priming buffer.

Cells were incubated in the pre-treatment solution for 35’ at room temperature, and digestion was quenched by adding 10 µL of 50 mg/mL bovine serum albumin (Fisher Scientific 12877172; dissolved in water). Cells were harvested by centrifugation at 1,250g for 5 minutes and resuspended in 25 µL SF buffer (Lonza) to a final concentration of 2E6 cells/µL. The resulting suspension is referred to as “primed cells”.

#### Nucleofection

For each individual nucleofection reaction, a nucleofection mix was prepared as follows: 4 µL SpCas9/gRNA RNP + 2 µL repair template (2 to 8 µg) + 2 µL primed cells (4E6 cells) + 10 µg pUC19 plasmid (used as carrier DNA) + SF buffer (Lonza) to a final volume of 24 µL. pUC19 plasmid was purchased from Tebu (066P-102:1mg), concentrated by ethanol/sodium acetate precipitation (see protocol in “Genomic DNA extraction” section below) and resuspended to ∼20 µg/µL.

For negative control reactions, RNP or repair template were replaced by the same volume of SF buffer. Cells were transferred into individual wells Lonza Nucleocuvette® 96-well plates or 16-well strips and nucleofected using the CM156 program for SF buffer. Immediately after nucleofection, 100 µL of ice-cold recovery buffer (10 mM HEPES-KOH pH. 7.5, 0.9 M sorbitol, 8% (weight/volume) PEG 8000) were added to each reaction. Cells were left to rest for 5’. Each 124 µL nucleofection reaction was split into two equal 62 µL aliquots (to obtain resistant clones from two independently transfected cells), each transferred in a separate well of a 6-well plate containing 2 mL of 1% (v/v) SWC in ASW. Cells were transferred into the 25°C culture incubator and incubated for 30 minutes before adding 0.5% (v/v) of a 10 mg/mL *E. pacifica* suspension in ASW.

### Puromycin selection

After 24 hours, 80 µg/mL puromycin (Sigma Aldrich P8833-10MG; dissolved in water to a stock concentration of 25 mg/mL and stored at −20°C) were added to each well.

### Clonal isolation

Clones were isolated by limiting dilution as in (Levin et al., 2014). After puromycin selection, resistant cells were diluted down to 0.5 cell/mL in 1% (v/v) SWC with 80 µg/mL puromycin and 1% (v/v) *E. pacifica* food pellet. For each transfected clone, two 96-well flat-bottom cell culture plates (ThermoFisher Scientific 130188) were filled with 200 µL diluted cell suspension per well. This corresponds to an expected value of 1 cell every 10 wells.

### Genotyping

Clonal isolation wells were monitored for cell growth. Clones were amplified for genotyping when cells reached high density (*i.e.* had visibly started to deplete food bacteria, but remained in chains and did not yet display the “fast swimmer” phenotype indicative of starvation (Dayel et al., 2011)). For each experiment, 6 clones were transferred into a 6-well plate (ThermoFisher Scientific 11825275) containing 3 mL ASW+1% (v/v) SWC with 80 µg/mL puromycin and 1% (v/v) *E. pacifica* food pellet. When cells reached high density in the 6-well plate, 1.5 mL of cell culture were centrifuged at 4,000g and the pellet was resuspended in 100 µL DNazol Direct (Euromedex 131) for genomic DNA extraction (following provider specifications).

Forward and reverse genotyping PCR primers were designed to flank the purported insertion site and be at least 50 bp distant from it (to allow Sanger sequencing with the same primers). Genotyping PCR reactions were set up with Q5 Polymerase following provider specifications, with 2-3 µL genomic DNA as template, 200 µM dNTP, 0.5 µM each primer and 0.02 U/µL Q5 polymerase.

The following PCR program was run: 30” 98°C; 40X (20” 98°C; 20” annealing temperature 68°C; 4’ 72°C); 4’ 72°C; hold 10°C. Annealing temperature was determined with the New England BioLabs Tm calculator: https://tmcalculator.neb.com/. A long elongation time (4 minutes) was chosen in order to detect potential tandem insertions of the puromycin resistance cassette in the target locus. Size of PCR products was assessed by electrophoresis on a 1% (w/v) agar gel and successful insertion of the puromycin resistance cassette was visualized as a 2 kb increase in PCR product size compared to the wild-type. For Sanger sequencing, PCR reactions were purified (PCR purification kit, Macherey NucleoSpin Gel and PCR cleanup 740609.50) and shipped to Eurofin Genomics.

### Rosette induction

Rosette development was induced by adding *Algoriphagus machipongonensis* bacteria. Frozen “food pellets” of *Algoriphagus machipongonensis* were prepared and stored at −80°C following an identical protocol to the one established for *E. pacifica* food pellets (Booth et al., 2018). Prior to use, each 10 mg food pellet was resuspended in 1 mL ASW and kept at 4°C. Rosette development was induced by adding 0.1% (v/v) *Algoriphagus* food pellet to cultures 16 to 24 hours prior to phenotype scoring.

### Whole-genome sequencing

#### Genomic DNA extraction

Genomic DNA was extracted using either DNazol (for the *rtls^pac1^* clone 1B5) or Qiagen Genomic DNA kit (Qiagen 13323).

DNazol extraction was performed as follows: ∼1E7 SrEpac cells were harvested by centrifuging ∼30 mL of a dense culture (∼3E5 cells/mL) for 15’ at 3,300 g, resuspended in 200 µL DNazol and transferred to a 1.5 mL Eppendorf tube. After 15’ room temperature incubation, DNA was precipitated by adding 0.1 volume sodium acetate 3M, mixing by pipetting up and down, 2.5 volume cold (−20°C) ethanol, mixing by vortexing, and letting precipitate overnight at −20°C. Samples were then centrifuged 30’ at 16,000g (maximal speed)/4°C, washed twice with ice-cold 70% ethanol:30% water (v/v), and centrifuged again for 15’ at 16,000g (maximal speed)/4°C. Supernatant was removed and pellet was air-dried for 5’ at 56°C before being resuspended in 50 µL EB buffer (from Macherey gel purification kit, reference 740609.50). DNA concentration was measured using a NanoDrop.

Qiagen Genomic DNA kit extraction was performed as follows: 50 mL of dense cell culture (>1E6 cells/mL) were concentrated into 2 mL 1% (v/v) SWC by 10’ 3,300g centrifugation, and left to graze bacteria overnight. Samples were then centrifuged 10’ 3,300g, supernatant was removed and pellet was stored at −20°C. DNA was then purified using the Qiagen Genomic DNA kit according to the provider’s protocol, resuspending the final pellet in 30 µL buffer NE. DNA concentration was measured using a NanoDrop. Samples were shipped to Eurofin Genomics for INVIEW Sequencing (Illumina 150 paired-ends sequencing).

#### Sequencing results analysis

Results were analyzed using Geneious Prime® 2024.0.4. Adapters were trimmed using BBDuk and reads were mapped onto the predicted *rtls^pac1^*reference genome (i.e. the *S. rosetta* reference genome downloaded from Ensembl Protists (Martin et al., 2023) with insertion of the termination/resistance cassette at the target site in the *rtls* locus). Mismatches were automatically annotated using the “Find variations/SNPs” tool with default parameters. In all *rtls^pac1^*clones, reads mapped to the termination/resistance cassette had homogeneous coverage (Figure S2) nearly identical to the average of the whole contig (GL832962) (Table S2), consistent with a single insertion. No mismatch was detected in reads mapping to the insert (except a one-base pair deletion at the beginning of the EFL promoter in clone 1B3; Figure S2, Table S2) consistent with a single specific insertion. By contrast, in no-RNP control clones, the coverage of the pEFL-pac-5’Act cassette was consistently 4-fold to 7-fold higher than the average of the contig (suggesting multiple copies per genome) and 70 to 80 mismatches were detected across the insert, notably at both extremities of the insert, clearly indicating presence of the cassette elsewhere than in the *rtls* locus. Inspection of mismatching reads indicated circularization of the cassette in both no-RNP control puromycin-resistant clones by recombination between the left and right homology arms (Figure S2) – and apparent episomal maintenance of the circularized fragment. Mismatches internal to the insert were polymorphic, suggesting differences between copies of the episomal element. Additionally, in one no-RNP clone only (clone 1D12), a significant number of internal reads were chimeric sequences between the repair template and the pUC19 plasmid co-transfected as carrier DNA, suggesting that recombination had taken place between the repair template and pUC19. Mapping of reads from this clone to the pUC19 reference sequence supported presence of the whole pUC19 plasmid, with coverage values suggesting 2 copies per cell on average. No read from any other clone (including the other no-RNP control) could be successfully mapped to pUC19, indicating that recombination of pUC19 with the puromycin resistance cassette and maintenance of the resulting chimeric plasmid was unique to this clone.

##### Growth curves

Growth curves were performed as in (Booth and King, 2020), in two independent replicates. For each KO genotype, two independent clones were assessed. Each clone was cultured in two independent flasks, and cell concentration in each flask was estimated from at least two independent cell density counts (measured with a LUNA-II^TM^ automated cell counter (LogosBiosystems)) at each timepoint. For wild type cells, two SrEpac cultures that had been passaged independently for more than 5 passages were thawed independently from a liquid nitrogen stock and were treated as independent “clones”. Doubling times *Dt* were quantified by comparing cell density *C(T)* during log phase (*T=*48 hours after seeding) to the initial cell density *C_0_* and computed with the following formula: *Dt* = *T*/[log_2_(*C(T)*/*C_0_*)].

##### Microscopy

###### Transmitted light imaging

For most experiments, *S. rosetta* cultures were imaged in untreated glass-bottom 96-well plates (Ibidi 89621), coated with 100 µL of 0.1 mg/mL poly-D-lysine (Sigma–Aldrich P6407-5MG) for about 30 seconds and rinsed twice with 100 µL distilled water. For rosette size quantification, rosette development was induced in standard conditions, in parallel for all genotypes. Briefly, cultures were first seeded at 5E4 cells/mL in T25 culture flasks (Fisher Scientific 11830765) in 10 mL culture medium supplemented with *Algoriphagus*: 5% (v/v) SWC, 1% (v/v) resuspended *E. pacifica* food pellet, 0.5 % (v/v) resuspended *A. machipongonensis* food pellet, 100 µg/mL kanamycin (Sigma Alrich K1377-1G; stored as a 50 mg/mL solution in water at −20°C), 100 µg/mL carbenicillin (Euromedex 1039-A; stored as 200 mg/mL in water), and 25 µg/mL tetracycline (dissolved to 12.5 mg/mL in a 1:1 water/ethanol solution and stored at −20°C). Antibiotics were included to ensure control and consistency of the bacterial communities during the experiment. Before passaging, mother cultures were briefly vortexed to dissociate chain colonies.

Rosette colonies were left to develop for 48 hours and were then collected for imaging as follows: 5 mL of each culture were harvested by centrifugation for 20’ at 5,000 g, resuspended in 1 mL ASW, purified with a Percoll solution as in (Levin and King, 2013) to reduce bacterial concentration, and resuspended in a final volume of 300 µL ASW. Cell suspensions were then transferred into an Ibidi 96-well plate that had been coated with poly-D-lysine. (For this experiment only, the poly-D-lysine solution was removed after coating without any additional washes with distilled water, as rosettes were found to require unwashed poly-D-lysine to be fully immobilized). Prior to imaging, a small quantity of paraformaldehyde (PFA) was added to immobilize rosettes (300 µL of a 1:2,000 dilution of a 16% PFA stock solution; Electron Microscopy Sciences 15714, to a final concentration, resulting in a final PFA concentration of 0.004%). Rosettes were imaged on a Zeiss Axio Observer 7 equipped with a Microscopy Camera Axiocam 712 mono (D) and a C-Apochromat 63x/1,20 W objective, using “Z stack” and “tile scan” options. Stacks were visualized using Fiji version 2.14.0/1.54f (Schindelin et al., 2012) and cells per rosette were manually counted for 50 rosettes per condition per biological replicate.

###### Confocal imaging

To image the 3D structure of wild-type and *warts^pac1^*mutant rosettes (Figure 5K,L), 300 µL of dense rosette cultures were first stained for ECM with fluorescein-jacalin (Eurobio Scientific FL-1151-5) and for membranes with FM 4-64 (Invitrogen T13320, resuspended according to provider’s specifications and added to a final working concentration of 1:1,000). Samples were then transferred into a 96-well plate coated with poly-D-lysine (see “Transmitted light imaging” above) and imaged live on a Leica Stellaris 5 confocal microscope with an HC PL APO 63x/1.20 W CORR CS2 objective and Lightning super-resolution mode.

## Supplementary Figures

**Figure S1.**
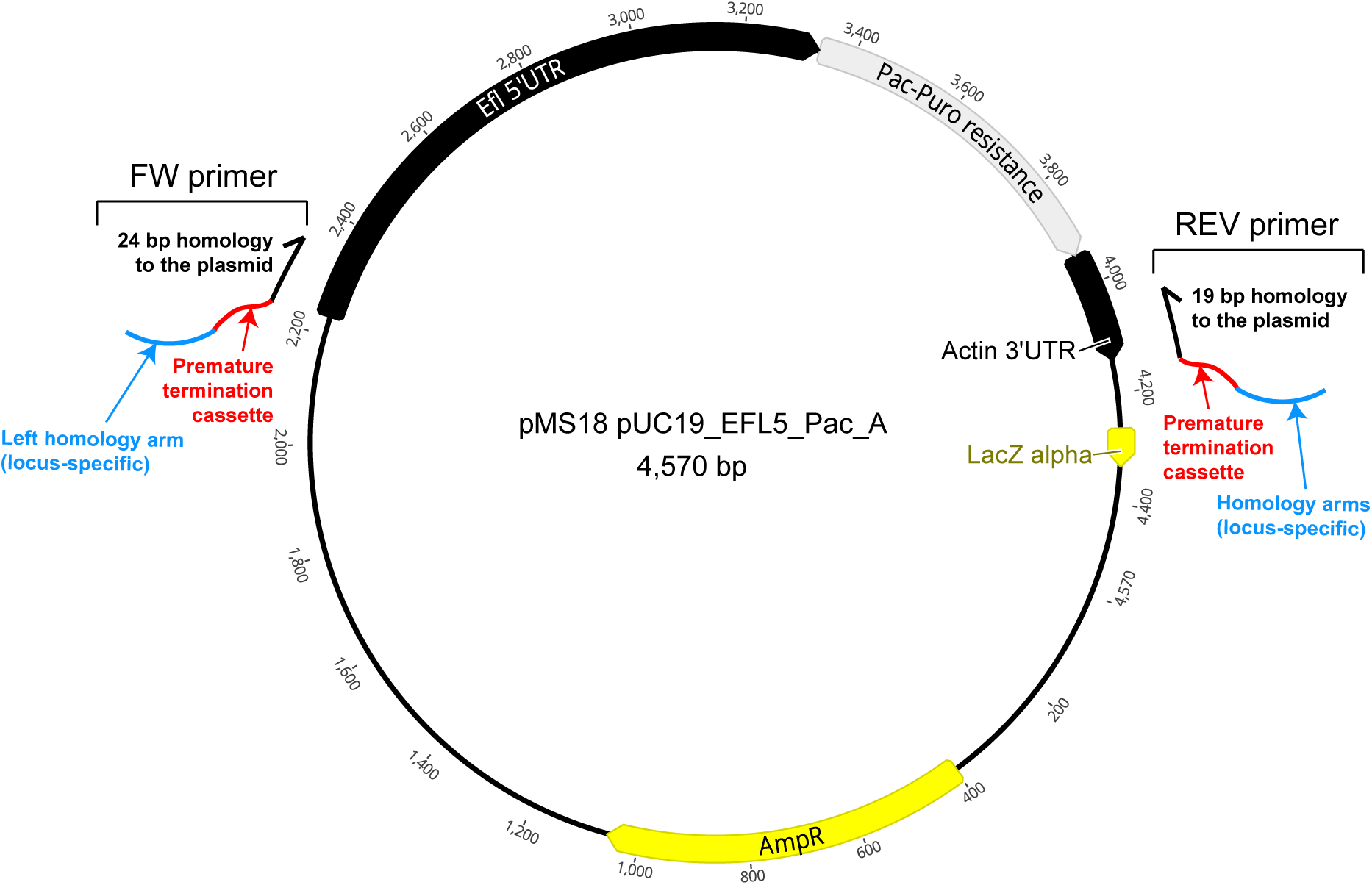
Repair template production pipeline. Repair templates are produced by PCR. A plasmid encoding a puromycin resistance cassette serves as a PCR template, and custom primers add premature termination sequences and homology arms to the genomic sequence flanking the Cas9 cutting site.

**Figure S2.**
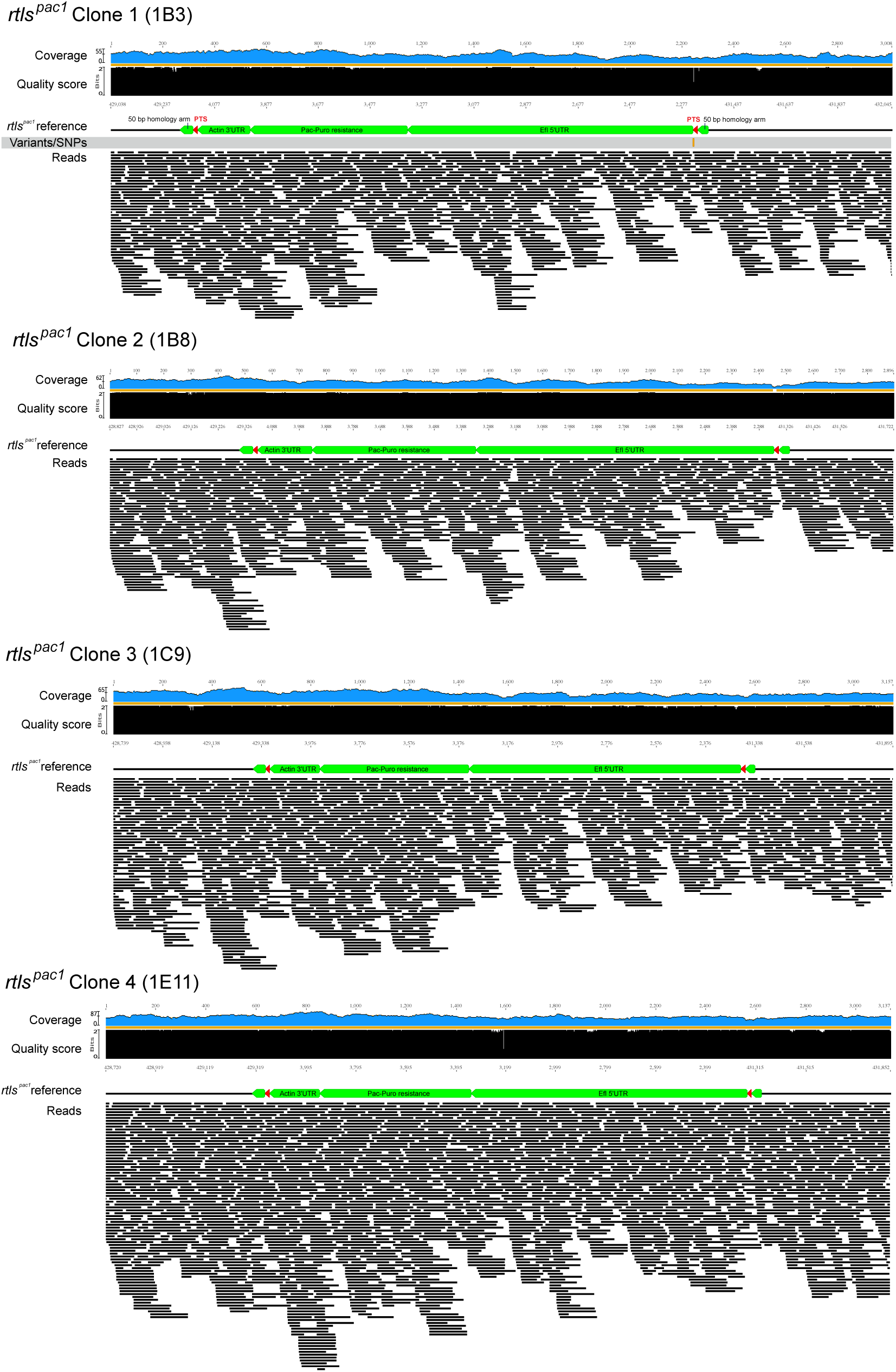
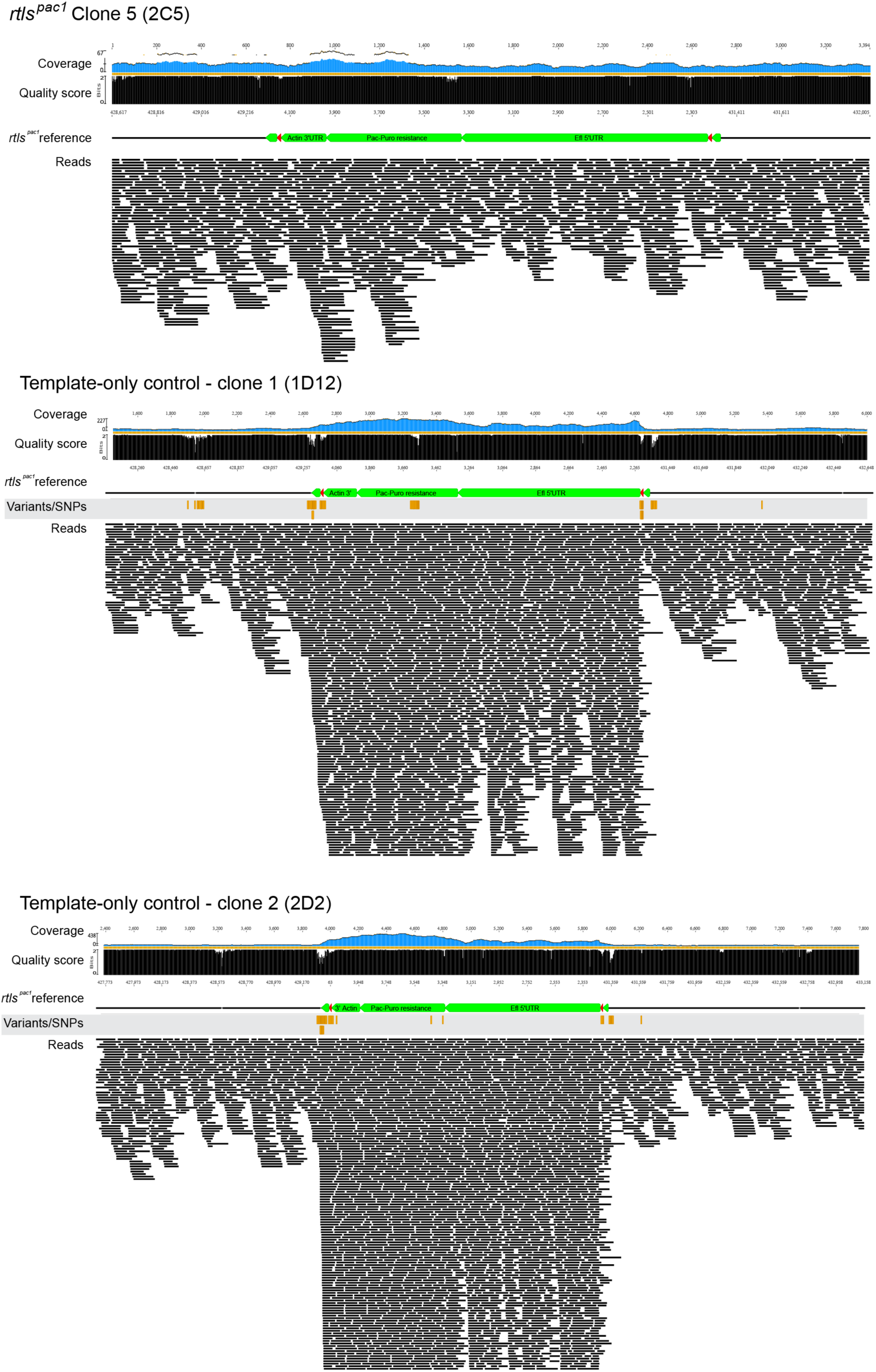
Whole-genome sequencing detects a single insertion of the termination/resistance cassette in five *rtls^pac1^* KO clones and detects multiple copies, all outside the *rtls* locus, in resistant template-only control clones. Reads mapping to the *rtls* locus of the reference *rtls^pac1^* genome in KO and template-only control clones. In *rtls^pac1^* clones, reads map to the *rtls* locus without any detectable deviation from the reference (“Variants/SNPs” line), except a deletion of one basepair in clone 1. Coverage of the termination/resistance cassette is similar to neighboring genomic regions, supporting a unique insertion. By contrast, in no-RNP (template-only) control clones, multiple mismatches are detected between reads and the predicted sequence, notably at both ends of the insertion site, consistent with the termination/resistance cassette not being inserted in the *rtls* locus. Moreover, coverage of the termination/resistance cassette appears much higher than that of the target locus, suggesting multiple copies per genome. See Table S1 for quantification.

**Table S1.** Statistics relating to whole-genome sequencing of *rtls^pac1^* knock-out and no-RNP negative control strains. A significant number of reads (red) could be mapped to the pUC19 plasmid (after recombination with the pEFL-pac-3’Actin cassette) in one no-RNP control clone only (1D12).

| Clone ID | Total number of reads | Number of reads mapped to pEFL-pac-3'Actin cassette | Number of reads mapped to pUC19 |
| --- | --- | --- | --- |
| *rtls<sup>pacI</sup>* clone 1 (1B3) | 24,065,034 | 562,378 | 4 |
| *rtls<sup>pacI</sup>* clone 2 (1B8) | 21,108,392 | 520,041 | 7 |
| *rtls<sup>pacI</sup>* clone 3 (1C9) | 21,160,812 | 609,234 | 0 |
| *rtls<sup>pacI</sup>* clone 4 (1E11) | 32,792,436 | 868,534 | 0 |
| *rtls<sup>pacI</sup>* clone 5 (2C5) | 21,150,366 | 636,365 | 0 |
| No-RNP control clone 1D12 | 24,297,494 | 16,386 | 1,708 |
| No-RNP control clone 2E2 | 17,815,344 | 572,990 | 0 |

**Figure S3.**
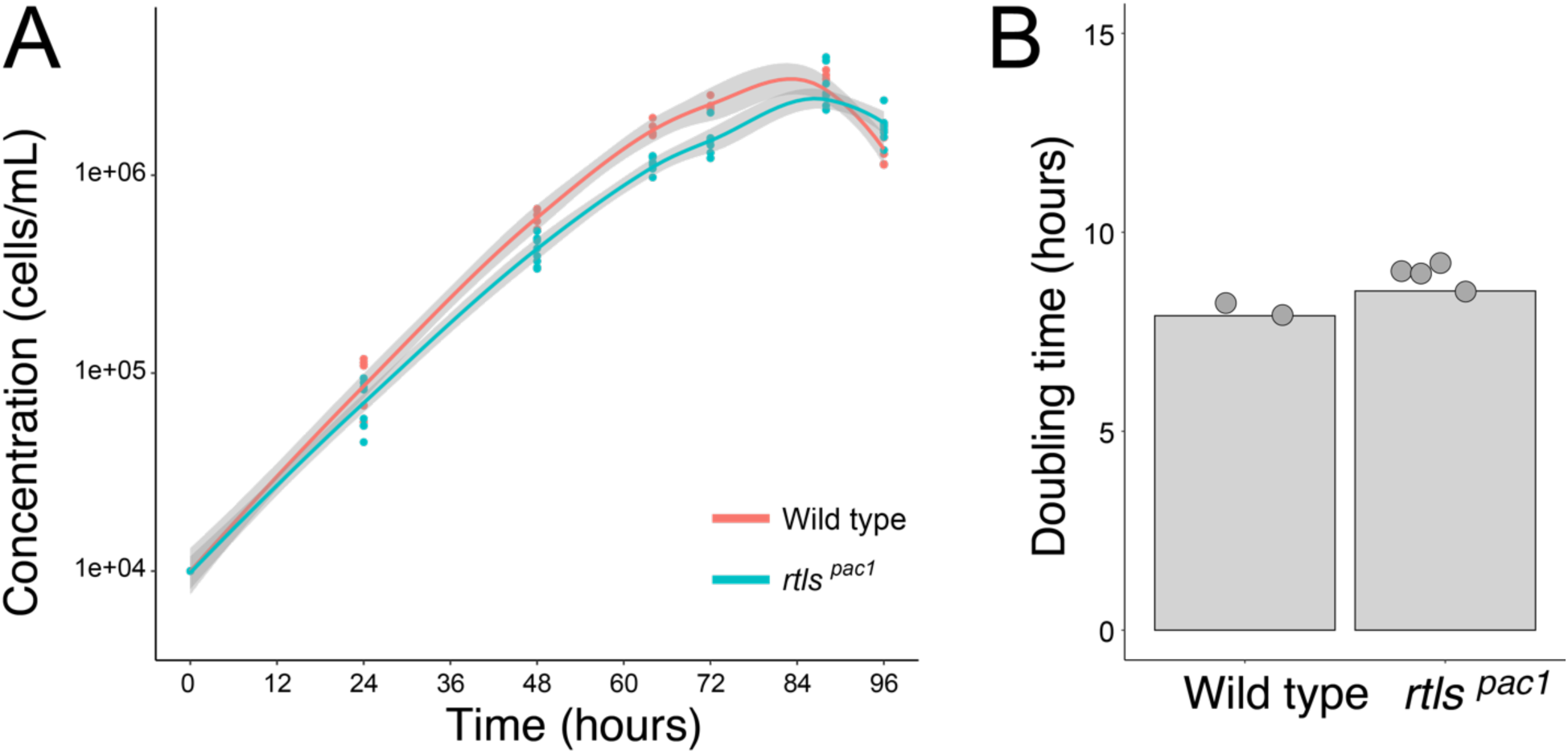
Wild type and *rtls^pac1^*KO cells grow at a similar rate. (A) Growth curves of wild type and *rtls^pac1^*cells (two distinct clones generated with 50-bp homology arms repair templates) at 25°C. (B) Doubling time did not significantly differ between wild type and *rtls^pac1^* cells (*p*=13.3% by the Mann-Whitney test).

**Figure S4.**
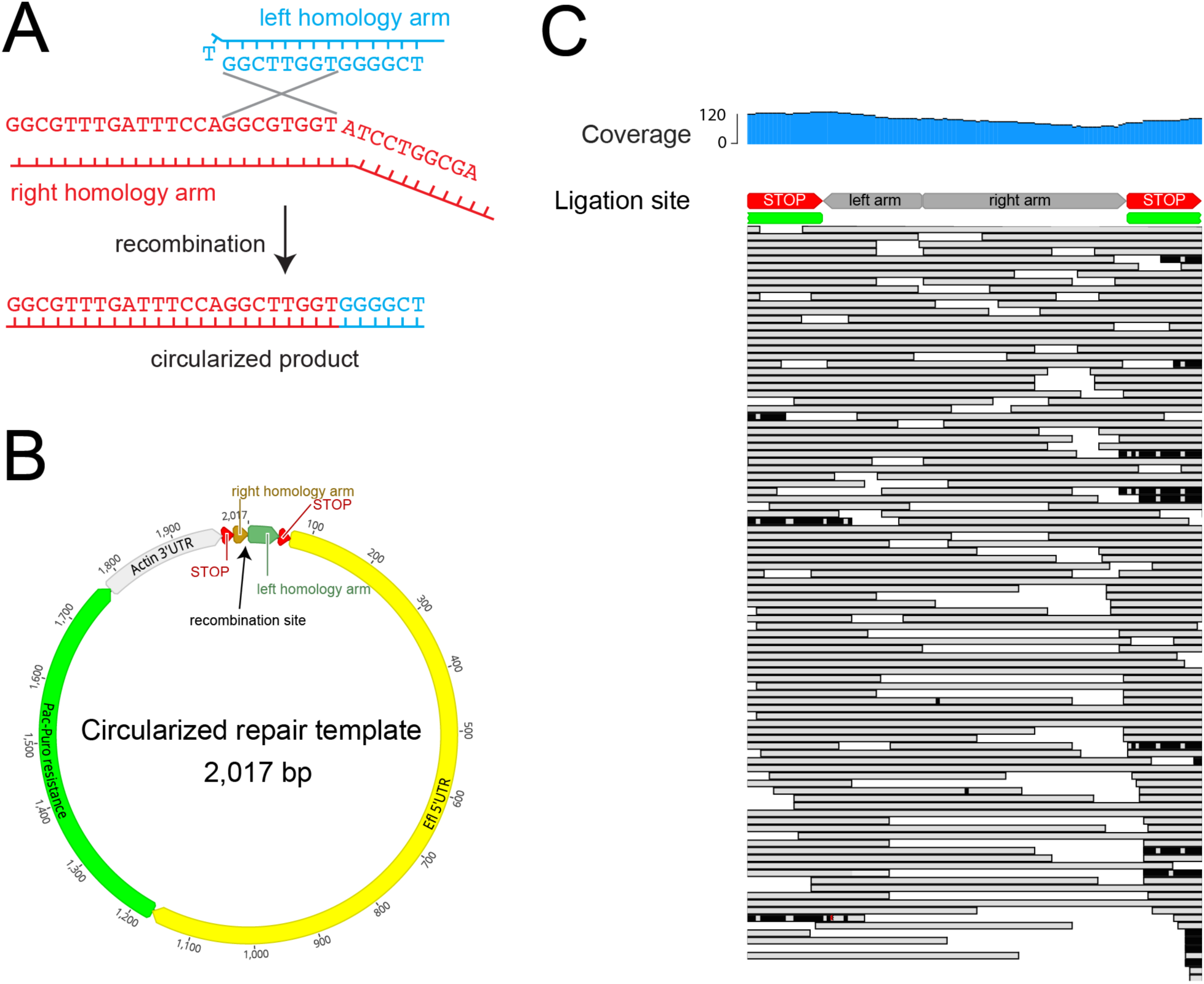
Repair template circularization and episomal maintenance can confer puromycin resistance in template-only control clones. (A) Recombination between the left and right homology arms of the *rtls* repair templates can give rise to a circular product. (B) Map of the circularized repair template produced by recombination. (C) Short-read Illumina sequencing reads aligning to the recombination site in puromycin-resistant template-only control clones. Multiple reads span the inferred recombination site, indicating presence of the circularized product.

**Figure S5.**
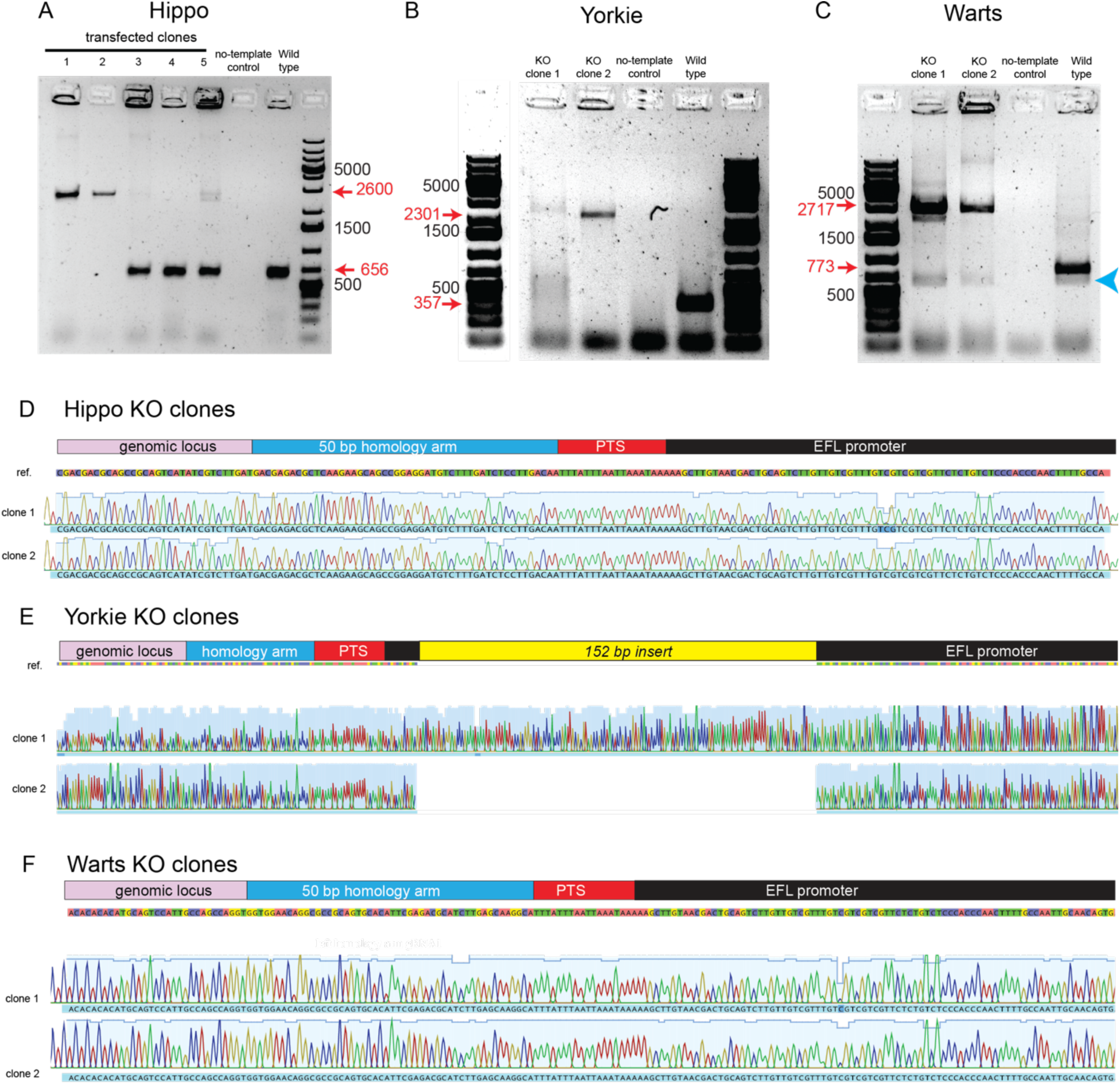
Genotyping of *hippo*, *warts* and *yorkie* KO clones. (A) Electrophoresis gel of genotyping PCR for *hippo^pac1^* clones. Five clones co-transfected with both RNP and repair template were genotyped, of which the first two (clone 1 and clone 2) displayed a unique band at the expected KO size (2,600 bp) and no band at the expected wild type size (656 bp). Only these two clones were considered KO and were characterized in further experiments. (B) Electrophoresis gel of genotyping PCR for *yorkie^pac^* clones. Both clone 1 and clone 2 lacked a band at the expected wild type size (357 bp) and displayed a band close to the expected KO size (2301 bp). Nevertheless, the band was slightly higher in clone 1 than in clone 2 and a PCR smear was also visible at lower sizes in clone 1 only, suggesting slightly different inserts between both KO clones, as was confirmed by sequencing (see panel E). We respectively named the clone 1 and clone 2 alleles *yorkie^pac2^* and *yorkie^pac1^*. The ladder (right) was overexposed compared to PCR products, and is duplicated and independently displayed with lower saturation on the left side of the gel image to allow better visual resolution of the ladder bands. (C) Electrophoresis gel of genotyping PCR for *warts^pac1^* clones. Both clone 1 and clone 2 lacked a band at the expected wild type size (773 bp) and displayed a band at the expected KO size (2717 bp). The gel imaged was the same one as in panel B (with the same ladder). The *hippo* and *yorkie* parts of the gels are shown in separate panels to adapt the contrast to the respective band intensities of the two sets of PCRs. An additional, non-specific PCR product (blue arrowhead), shorter than the wild-type *warts* product, was present in all reactions. (D-F) Sanger sequencing confirmed insertion of the termination/resistance cassette in *hippo^pac1^*, *yorkie^pac^* and *warts^pac1^*clones. In *yorkie^pac2^* (*yorkie* clone 1), an additional, unexpected 152 bp insert was detected close to the 5’ end of the EFL promoter. The insert is a tandem repeat of the previous 152 bp within the locus (including part of the EFL promoter, the termination sequence, the left homology arm, and 62 bp of the genomic context). A second, similar tandem repeat (of 173 bp) was also detected at the 3’ end of the PCR product, covering part of the Actin 3’ UTR, the termination sequence, and 34 bp of the right homology arm. This suggests that, in clone 1 (*yorkie^pac2^*) only, tandem repeats were inserted at both 5’ and 3’ ends by the DNA repair machinery, explaining the slightly larger insert size (∼300 bp larger) observed by electrophoresis as well as the smear at lower sizes (likely due to repeated sequences causing incomplete PCR products of variable length). As the *yorkie* locus had been properly disrupted, and although the inserted sequence was slightly different than initially intended, the *yorkie^pac2^* clone 1 was considered a *bona fide* KO and was investigated alongside *yorkie^pac1^* clone 2 in downstream experiments.

**Figure S6.**
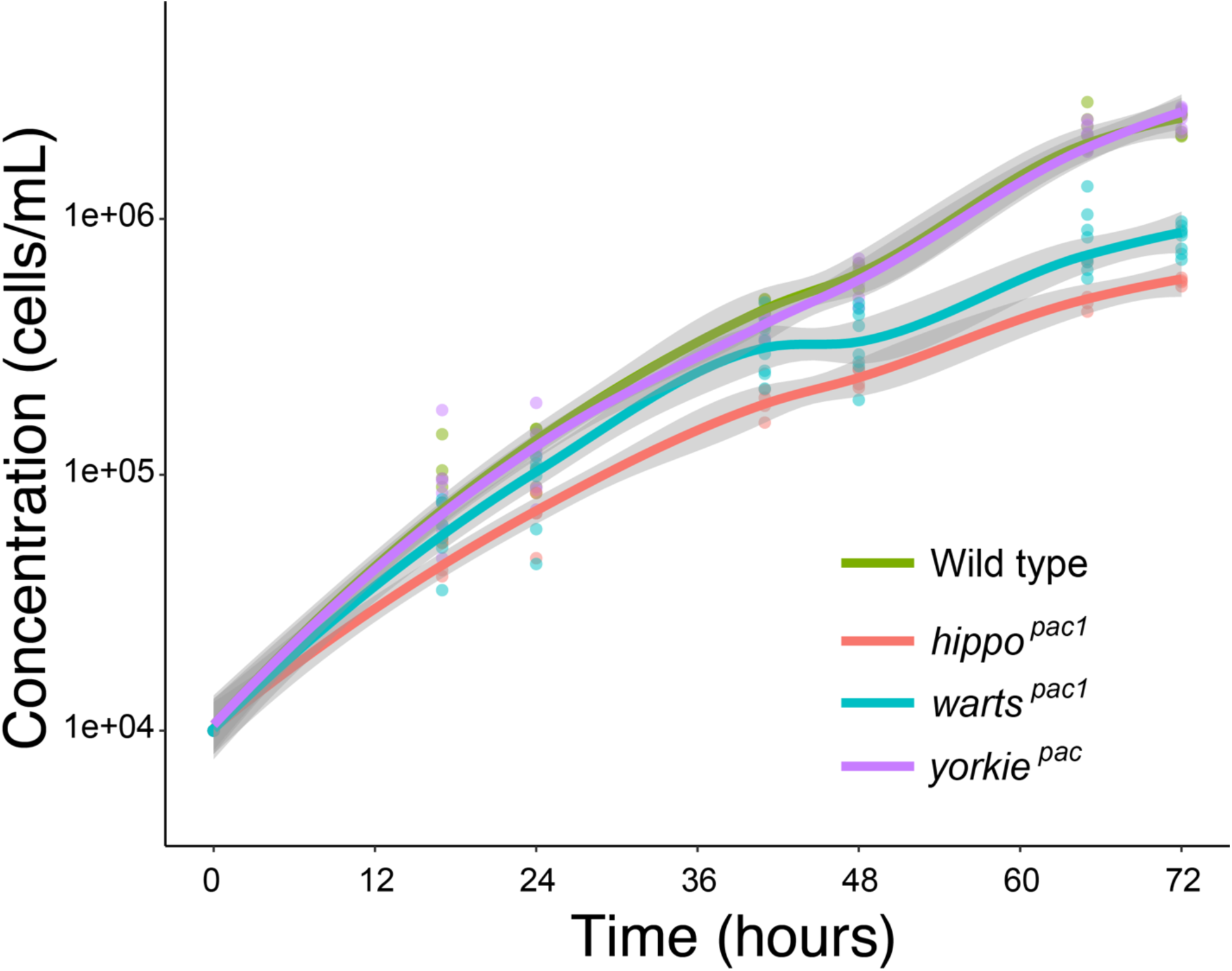
Independent biological replicate for growth curves of wild type *S. rosetta* compared to Hippo pathway KO strains. Same experimental procedure as in Fig. 5B. Similar differences in growth rates between strains were observed. Doubling times quantified in this experiment are plotted in Fig. 5C (“Replicate 2”).

